# Vestibular afferent neurons develop normally in the absence of quantal/glutamatergic input

**DOI:** 10.1101/2024.06.12.597464

**Authors:** Katherine Regalado Núñez, Daniel Bronson, Ryan Chang, Radha Kalluri

**Affiliations:** Neuroscience Graduate Program, University of Southern California, Los Angeles, CA, 90089; Caruso Department of Otolaryngology-Head and Neck Surgery, Keck School of Medicine, University of Southern California, Los Angeles, CA, 90033; Zilkha Neurogenetic Institute, Department of Otolaryngology, University of Southern California, Los Angeles, CA, 90033; Dornsife College of Letters, Arts and Sciences, University of Southern California, Los Angeles CA 90089

**Keywords:** Vestibular, VGLUT3, KCNQ, Afferent development, Afferent diversity. (Min.5- Max. 8)

## Abstract

The vestibular nerve is comprised of neuron sub-groups with diverse functions related to their intrinsic biophysical properties. This diversity is partly due to differences in the types and numbers of low-voltage-gated potassium channels found in the neurons’ membranes. Expression for some low-voltage gated ion channels like KCNQ4 is upregulated during early post-natal development; suggesting that ion channel composition and neuronal diversity may be shaped by hair cell activity. This idea is consistent with recent work showing that glutamatergic input from hair cells is necessary for the normal diversification auditory neurons. To test if biophysical diversity is similarly dependent on glutamatergic input in vestibular neurons, we examined the maturation of the vestibular epithelium and ganglion neurons in *Vglut3-ko* mice whose hair cell synapses lack glutamate. Despite lacking glutamatergic input, the knockout mice showed no notable balance deficits and crossed challenging balance beams with little difficulty. Immunolabeling of the *Vglut3-ko* vestibular epithelia showed normal development as indicated by an identifiable striolar zone with calyceal terminals labeled by molecular marker calretinin, and normal expression of KCNQ4 by the end of the second post-natal week. We found similar numbers of Type I and Type II hair cells in the knockout and wildtype animals, regardless of epithelial zone. Thus, the presumably quiescent Type II hair cells are not cleared from the epithelium. Patch-clamp recordings showed that biophysical diversity of vestibular ganglion neurons in the *Vglut3-ko* mice is comparable to that found in wildtype controls, with a similar range firing patterns at both immature and juvenile ages. However, our results suggest a subtle biophysical alteration to the largest ganglion cells (putative somata of central zone afferents); those in the knockout had smaller net conductance and were more excitable than those in the wild type. Thus, unlike in the auditory nerve, glutamatergic signaling is unnecessary for producing biophysical diversity in vestibular ganglion neurons. And yet, because the input signals from vestibular hair cells are complex and not solely reliant on quantal release of glutamate, whether diversity of vestibular ganglion neurons is simply hardwired or regulated by a more complex set of input signals remains to be determined.

## 1 Introduction

Neural signal processing is supported by biophysically diverse neurons that respond to specific qualities of the input stimuli. Vestibular ganglion neurons (VGN, the cell bodies of vestibular afferent nerve) are a remarkable example of a single neuron population with a wide range of biophysical diversity. A subset of VGN containing large densities of low-voltage gated potassium channels (Kv1 and KCNQ) are signal differentiators producing action potentials only at the onset of current step injections (so called ‘transient-firing’ neurons), whereas others having little to no low-voltage gated potassium channels fire long repetitive spike trains in response to the integrated charge of the incoming signal (so-called ‘sustained-firing’ neurons) (Iwasaki et al., 2008; Kalluri et al., 2010). Such diversity in the intrinsic biophysical properties of vestibular neurons is believed to influence their different functions. VGN with transient-firing are associated with irregular-firing afferents (Kalluri et al., 2010) that predominantly innervate the central zones of vestibular epithelia (Goldberg, 2000) and are suited for encoding rapid/dynamic changes in head position through the precise timing of their individual spikes (Sadeghi et al., 2007). In contrast, VGN with sustained-firing are associated with regular-firing vestibular afferents whose action potentials come at uniform intervals, predominantly innervate the peripheral regions of vestibular epithelia, and are believed encode slow changes in head position via their average rate of spiking. Here, we ask if the intrinsic biophysical diversity of vestibular ganglion neurons is shaped by activity dependent regulation of ion channels during post-natal development.

We hypothesized that removing/reducing the activity from hair cells to vestibular afferents during post-natal development will reduce biophysical diversity while also impairing vestibular function. This hypothesis is motivated in part by recent observations in auditory neurons showing that the diversification of spiral ganglion neurons (SGN) depends on receiving input from inner hair cells during development. Specifically, SGN of vesicular glutamate transport 3 knockout mice (*Vglut3-ko*), which lack glutamatergic release, fail to diversify into three molecular subtypes defined by single cell sequencing of adult SGN (Petitpré et al., 2018; Shrestha et al., 2018; Sun et al., 2018). Lack of diversification is marked by a failure to upregulate the expression of potassium channels and a failure to refine the expression of key molecular markers of SGN subtype such as the calcium-binding protein calretinin. Consistent with the failure to upregulate potassium channels, patch-clamp recordings found that SGN in the *Vglut3-ko* mice are hyper-excitable when compared to those from wild-type mice (Babola et al., 2018). While molecular and biophysical diversity is already evident in SGN before hair cells are fully transducing (Petitpré et al., 2018; Markowitz and Kalluri, 2020), functional diversity in the adult system may be driven by a combination of genetic hardwiring and post-natal maturation. Hyper-excitability in the KO mice may be a juvenile phenotype when potassium channels fail to developmentally upregulate. Overall, the data suggest that ion channel expression (specifically potassium channels) in SGN is regulated by the sensory input received from inner hair cells during post-natal development. Thus activity-dependent maturation is an important component of functional diversification in the spiral ganglion neuron and may similarly be important for the vestibular ganglion neurons.

We employed the *Vglut3-ko* mouse as the experimental model because the hair cell synapses in these mice are missing glutamate (Seal et al., 2008). Typically, deflections of vestibular hair cell bundles caused by head movements lead to the depolarization of hair cells and an influx of K^+^ and Ca^2+^, the latter of which facilitates the quantal release of glutamate into the synaptic cleft. The released glutamate elicits excitatory post synaptic currents that trigger vestibular neurons to fire action potentials (Eatock and Songer, 2011). Glutamate is the primary neurotransmitter released by vestibular hair cells since pharmacologically blocking post-synaptic glutamate receptors quenches the excitatory post-synaptic quantal currents in wildtype mice (Bonsacquet et al., 2006; Sadeghi et al., 2014). Multiple lines of evidence suggest that blocking glutamate release will profoundly reduce the input that vestibular afferents receive during development. All hair cells in the vestibular epithelium contain multiple active zones containing glutamate-bearing synaptic ribbons (Lysakowski and Goldberg, 2008), thus significant energy is used to maintain this synaptic machinery. Based on immunohistochemistry and single-cell sequencing in neonatal mice, VGLUT3 is the exclusive glutamate transporter supplying hair cell ribbon synapses in both mouse and zebra fish hair cells (Wang et al., 2007; Obholzer et al., 2008; Seal et al., 2008). Neither VGLUT1 nor VGLUT2 compensate for the loss of VGLUT3 in zebra fish (Obholzer et al., 2008). While glutamate transporters like EEAT4/EEAT5 are found in mouse vestibular hair cells (Dalet et al., 2012), this class of transporter is typically shuttling away released glutamate rather than filling vesicles. Thus, the weight of existing data supports the idea that neither auditory nor vestibular hair cells release glutamate in *Vglut3-ko* mice.

Unlike in the auditory system where the *Vglut3-ko* produces complete knockout of inputs from the hair cell, the scenario is more complex in the vestibular system. Vestibular epithelia have two types of hair cells, one of which can communicate by both quantal release of glutamate and via a non-quantal mechanism that is not dependent on glutamate. Type I hair cells are paired with a unique cup-shaped post-synaptic terminal that envelops the whole basal pole of the hair cells (calyx terminal). This unique geometry facilitates non-quantal transmission which is proposed to be driven by a variety of mechanisms including in part by the accumulation of K^+^ in the post-synaptic cleft upon release from Type I hair cells, elevated electrical potential and/or acidification in the narrow cleft, and resistive coupling between low-voltage gated ion channels found in the cleft facing membranes of the hair cell and calyx terminal (Yamashita et al., 1990; Goldberg, 1996; Eatock and Songer, 2011; Highstein et al., 2014; Contini et al., 2020; Govindaraju et al., 2023). Still, the exact mechanism and the degree to which nonquantal transmission contributes to synaptic transmission between vestibular hair cells and vestibular afferents has yet to be determined.

We tested vestibular function in *Vglut3-ko* mice at two and three months of age by comparing balance beam performance to age-matched control mice. We tested if the regional organization of vestibular epithelia was impacted in the knockout mouse by looking for the developmental acquisition of calretinin and KCNQ4 ion channels in the calyx-only endings of central-zone afferent nerve endings. We also quantified the number and proportion of Type I and Type II hair cells. We were especially interested in asking if the putatively silent Type II hair cells were maintained into maturity or pruned away. Finally, we used whole-cell patch clamp recordings from VGN somata to compare the biophysical diversity of VGN in wild-type and *Vglut3-ko* mice in first, second, and third post-natal weeks.

We report that despite lacking quantal transmission, the biophysical properties, epithelial organization, and vestibular function of the *Vglut3-ko* mice appear normal when compared to controls. Functional tests in *Vglut3-ko* mice show normal vestibular performance on balance beam tests. Epithelial morphology in the *Vglut3-ko* was largely intact, and a quantification of hair cells in three areas of the utricular macula found no significant difference in the number or type of hair cells present compared to controls. Whole cell recordings of vestibular afferents in *Vglut3-ko* were appropriately diverse in their firing patterns, with both transient and sustained-firing populations present at the end of the first, second, and third postnatal week. Despite largely normal vestibular behavior and development, we observed a slight but significant increase in excitability in large, putatively central zone projecting VGN.

## 2 Methods

### Animal Model and Preparation

Data were recorded from the superior portion of the vestibular ganglion of mice from the C57BL6/J background strain of either sex aged post-natal day (P)8-25 (P0, birth day). Breeding pairs for homozygous *Vglut3-ko* mutant mice were provided by J. Oghalai and were originally received from the Jackson Laboratory (RRID:IMSR_JAX:016931). All animals were handled and housed in accordance with National Institutes of Health *Guide for the Care and Use of Laboratory Animals.* All animal procedures were approved by the University of Southern California Institutional Animal Care and Use Committee.

Dissection, dissociation, and culturing of vestibular afferent neurons follows procedures that we have previously described in detail (Kalluri et al., 2010; Iyer et al., 2023), and is summarized as follows. The superior ganglion contains the cell bodies of the afferents that project from the utricle, as well as the lateral and horizontal cristae. The inferior ganglion was explicitly excluded from this study since we’ve observed that dissociation is more difficult when it is included. Chemicals were obtained from Sigma-Aldrich unless otherwise specified. Temporal bones were dissected from the animals in chilled and oxygenated Leibowitz medium supplemented with 10 mM HEPES (L-15 solution). The superior part of the vestibular ganglia was detached from the distal and central nerve branches and separated from the otic capsule. Bone fragments, debris and any remaining connective tissue were removed from the surface of the ganglia. 2 ganglia from 1 to 2 litter-matched animals of either sex were pooled together. Ganglia were then incubated at 37°C in L-15 solution with 0.05% collagenase, 0.25% trypsin and 0.05% DNAse I for 20-40 min, depending on age of the animal. DNAse I was added to the enzymatic solution in this study to increase cell survival and reduce clumping of cells during trituration. We chose this enzymatic cocktail since it appears to preserve cell size and age-dependent ion channel function that is consistent with other methods that don’t rely on enzymatic treatment (Risner and Holt, 2006; Rocha-Sanchez et al., 2007; Kalluri et al., 2010; Meredith and Rennie, 2015; Ventura and Kalluri, 2019).

After enzymatic treatment, ganglia were washed sequentially in fresh L-15 solution, and culture medium (see below). Somata were dissociated in culture medium by triturating through a series of polished Pasteur pipettes and allowed to settle onto poly-D-lysine coated coverslips (MatTek Corporation). Culture dishes contained bicarbonate-buffered culture medium (minimal essential medium, Invitrogen), supplemented with 10 mM HEPES, 5% FBS, and 1% penicillin streptomycin (Invitrogen). The culture medium was titrated with NaOH to a pH of 7.35. Cells were incubated for 16-24 hours in 5% CO_2_/95% air at 37°C. Short-term incubation tends to remove supporting and satellite cells, clears debris from enzyme treatment, allows cells to adhere to the substrate, and promotes successful recordings as the animals entered the second post-natal week and beyond.

### Electrophysiology

Cells were viewed at 400x using an inverted microscope (Zeiss, Axiovert 135 TV) fitted with Nomarski optics. A MultiClamp 700B amplifier, Digidata 1440 board, and pClamp 10.7 software (MDS; RRID: SCR_011323) were used to deliver, record, and amplify all signals. Recording pipettes were fabricated using filamented borosilicate glass. Pipettes were fire polished to yield an access resistance between 4 and 8 MΩ. Recording pipettes were coated with parafilm (Bemis Company Inc) to reduce pipette capacitance.

Patch clamping was performed using the perforated-patch method. The contents of the perforated-patch internal solution contained the following (in mM): 75 K_2_SO_4_ ,25 KCl, 5MgCl_2_, 5 HEPES, 5 EGTA, 0.1 CaCl_2_, and titrated with 1 M KOH to a pH of 7.4 and an osmolality of 260 to 280 mmol/kg. Amphotericin B (240/ml; Sigma-Aldrich) was dissolved in DMSO and added to the perforated-patch solution on day of recording. This allowed passage of small monovalent ions while preventing larger molecules from dialyzing.

The series resistance was estimated in voltage-clamp using the built-in series resistance estimation function on the MultiClamp. Series resistance ranged between 8 and 35 MΩ. Recordings with higher than 35 MΩ series resistance were excluded from analysis. The same cutoff value of 35 MΩ has been used in our previous studies (Ventura and Kalluri, 2019; Bronson and Kalluri, 2023). Series resistance was left uncompensated. Although we recorded series resistance to ensure proper access, we used uncorrected voltage-clamp data due to the challenges of obtaining stable measurement of series resistance when using perforated-patch technique. Electrophysiology recordings show uncorrected command voltages as indicated on the figures.

Whole-cell capacitance was estimated online using the built-in estimation circuitry on the MultiClamp amplifier. On-line calculation of capacitance (*C*_m_) was verified off-line by analyzing recordings of capacitive currents in response to small depolarizing voltages. Offline verification was done by first fitting a single exponential to the transient current evoked by a 5-mV depolarizing step and measuring the membrane time constant (*τ*_m_). *C*_m_ was calculated by dividing *τ_m_* by the series resistance. Input resistance (*R*_in_) was calculated from voltage changes in response to a hyperpolarizing 10 or 20 pA step in current-clamp mode.

Recordings were made at room temperature (25-27°C) and in an external bath continuously perfused with fresh oxygenated L-15 media. Perforated patch internal solution had junction potentials of +5.0 mV, which was computed with JPCalc (Barry, 1994) as implemented by pClamp 10.7 and left uncorrected. Only recordings in which the cell had formed a giga-ohm seal were used. Recording stability was monitored by measuring resting potential, input resistance, series resistance and size of action potential. Significant fluctuations in these values within each experimental condition was indicative of an unhealthy cell and/or compromised recording and was not included in our dataset.

### Electrophysiology Analysis

All data were analyzed with pClamp 10 software (Clampfit; MDS Analytical Technologies) and Matlab R2022b (Mathworks). An automated Matlab script was used to generate counts of the number of action potentials in traces recorded in current clamp mode. Action potentials were identified when the rate of voltage increase exceeded an exponential increase of 5 millivolts per millisecond. A smoothing function was used to ensure that the script did not mistakenly count duplicate action potentials. After the script flagged individual spikes, each trace was verified manually to ensure the correct number of spikes were identified.

### Electrophysiology Statistics

Statistical analysis was done with Origin Pro (OriginLab; RRID: SCR_014212) and/or JMP Pro 13 (SAS Institute; RRID:SCR_014242). Repeated measures and two-factor ANOVAs were used to assess statistical significance when comparing between experimental and groups across multiple levels. In this case, post-hoc differences were estimated with Tukey’s HSD to avoid family-wise error. Student t-tests were used when comparing only two groups. Assumptions of equal variance and normality were assessed prior to running t-tests with Levene’s and Kolmogorov-Smirnov tests, respectively.

### Immunohistochemistry

Otic capsules were extracted from euthanized mice aged P15 and P31, and fixed in 4% paraformaldehyde for 1 hour, followed by three consecutive washes in phosphate-buffered saline (PBS). Then, vestibular epithelial tissue was extracted while the otic capsule was submerged in PBS. The extracted tissue was incubated at room temperature for one hour in a blocking solution containing goat serum dilution buffer (GSDB) with 4% Triton X. The blocking solution was then washed off with one round of PBS. Next, the tissue was incubated between 16 and 24 hours at room temperature in a primary antibody dilution containing GSDB with 0.3% Triton X. After incubation, the primary solution was washed off with three rounds of PBS at ten minutes per wash. The tissue was then incubated at room temperature for one hour in a solution containing secondary antibodies diluted in GSDB with 0.3% Triton X. Finally, tissue was washed with three rounds of PBS at ten minutes per wash, and mounted.

The primary antibodies and corresponding dilutions used are as follows: (1) anti-CtBP2 mouse monoclonal antibody IgG1 (BD Transduction Laboratories) at 1:200, (2) anti-Calretinin rabbit polyclonal antibody (Swant) at 1:1,000, (3) anti-b-tubulin III mouse monoclonal antibody IgG2a (Covance) at 1:200, (4) anti-KCNQ4 mouse monoclonal antibody IgG1 (NeuroMab) at 1:400. The secondary antibodies were used, all in a dilution of 1:200: Alexa Fluor 633 anti-rabbit (Molecular Probes), Alexa Fluor 594 anti-mouse IgG2A (Molecular Probes), Alexa Fluor 488 anti-mouse IgG1 (Life Technologies), and Alexa Fluor 647 anti-mouse IgG1 (Invitrogen). Note that the immunostaining for KCNQ4 was run separately from the rest of the immunostaining, and on different tissue, with secondary Alexa Fluor 647 anti-mouse IgG1.

### Imaging

Prior to mounting, clear nail polish was used to make four dots on the glass slides, creating a raised surface for glass cover slips to sit on. Immunolabeled slides were then mounted on the glass slides using mounting medium Vecta-Shield (Vector Laboratories). Mounted tissue was imaged using a Zeiss LSM 800 confocal microscope with a 20x lens at 0.5x magnification, and a 63x oil-immersion lens at 1x magnification. We captured z-stacks that were 512x512 pixels, with 2x averaging set to Sum mode to counteract dimness of some of the scans. Z-stacks were obtained with z-steps of optimal sizes per lens (0.56 µm for 20x lens, 0.24µm for 63x oil lens). To account for variation between samples, laser power and gain was adjusted accordingly per scan.

### Hair Cell Count

Hair cells were identified and quantified using 63x z-stacks of whole mount utricles from three C57BL6J mice and three Vglut3KO mice, all aged P15. Scan dimensions were 101.41µm by 101.4µm. Three regions of the utricle were inspected: the medial rostral extrastriola, medial caudal extrastriola, and central utricle (encompassing the striola and juxtastriola). We immunolabeled with CtBP2, calretinin, and β-III Tubulin. To identify the vestibular hair cells, we used nuclear stain CtBP2 and parsed through utricle scans. While CtBP2 also labels supporting cells in vestibular epithelia, supporting cell nuclei can be distinguished as they are found lower in the epithelium and have irregularly shaped nuclei (Fernandez et al., 1995). Once the layer of nuclei belonging to hair cells was identified, we reviewed the colocalization of CtBP2 and neuronal marker β-III Tubulin. As β-III Tubulin labels vestibular afferent calices, CtBP2-positive nuclei located within a β-III Tubulin-labeled calyx were identified as belonging to a Type I hair cell. The remaining CtBP2-labeled nuclei at the hair cell layer were identified as Type II hair cells. The majority of these had colocalization with a calretinin-labeled hair cell. Calretinin primarily labels Type II hair cells in the extrastriolar zone of the utricle, and calices in the striola (Desai et al., 2005). If a nucleus was found within a β-III Tubulin-labeled calyx and also colocalized with calretinin, the nucleus was identified as a Type I hair cell, as calretinin has been shown to label a minority of Type I hair cells (Desai et al., 2005). Hair cell identification was reviewed with orthogonal views of utricles.

### Balance Testing

A balance beam test was selected as it is a useful measure of balance and motor coordination. A previous study used beam-crossing tests to characterize balance and motor coordination in transgenic mice with known deficits in only one zone of the vestibular epithelium (Ono et al., 2020).

A balance beam apparatus was constructed with its beam elevated 50 cm from the base to dissuade mice from attempting to leave the trial. To encourage mice to complete the trial, a goal box was placed at the end of the beam with nesting material inside. A nylon hammock was suspended below the beam to prevent injury in case of falls. Balance beams of 12 mm and 6 mm in width were used in this experiment. The 12 mm beam provided a more easily accomplished task for the mice, while the 6 mm beam provided a more difficult task. Beam traversal times were compared at each beam width between the two mouse strains. Mice aged P15-P18, P30-P40 and P70-P80 were included. Animals in the P15-P18 age range were not able to complete the task. (No age dependent trends were noted between the two older groups, and the data was thus pooled).

Prior to the experiment, mice were placed in the procedure room for 15 minutes to acclimate to the environment. Here we defined a trial as the attempted traversal by a mouse across an 80 cm distance towards the goal box. Before the execution of individual trials, mice are placed inside of the goal box for 15 seconds. Placement of the mice in the goal box is intended to provide motivation for their return once placed on the opposite end of the beam. During the trial, a mouse is placed on one end of the beam 100 cm away from the goal box. The middle 80 cm of the traversal is used for data collection and analysis. The time of traversal, the number slips, the number of stalls, and the number of prods that occurred during a mouse’s 80 cm traversal were recorded. A slip is defined as a significant portion of the mouse’s body falling below the surface of the balance beam or a limb sliding below the bottom of the beam. A stall is defined as a significant interruption in traversal of the mouse. A prod is a gentle poke to the backside of the mouse to re-motivate traversal if a stall persists for an extended period (therefore all prods include stalls).

On a trial day, mice first complete two trials on the 12mm beam with 15 seconds between trials to conserve the motivation for a mouse to complete the trial and to provide the mice with a short break. After the completion of two trials, mice are placed back inside of their home cage to rest for 10 minutes. After resting for 10 minutes, mice complete two trials on the 6mm beam using the same procedure as the 12mm beam. During this resting period, other mice were tested. Each mouse was tested on two days (with one day in between) for a total of 4 trials per mouse on each beam.

## 3 Results

We evaluated the maturation and organization of the vestibular epithelium in *Vglut3-ko* mice in comparison to wild-type mice using the following: 1) balance beam tests to identify potential large-scale changes in balance and motor function, 2) immunohistochemistry to examine the regional organization of the utricle and cristae of the semicircular canals with an emphasis on features that are typically different between central and peripheral zone afferents and 3) electrophysiological characterization of neuronal diversity using patch-clamp recordings from vestibular ganglion neuron soma.

### 3.1. Knockout animals are able to cross balance beams with no obvious deficits

Gross vestibular function is largely intact in *Vglut3-ko* mice (see Methods for detailed protocol). Mice had normal righting reflex in early post-natal days and did not show obvious defects in their cage behavior. We evaluated motor behavior with a balance beam crossing paradigm, which is a typically challenging task that has previously been effective at revealing peripheral deficits (Ono et al., 2020). Mice were required to cross an 80 mm spans of 12 mm and 6 mm elevated balance beam. Balance beam traversal time, slips and stalls were measured and compared between wildtype mice (n=11) and *Vglut3-ko* mice (n=13) from both juvenile (P30-P35) and adult animals (P70-75). Each animal crossed a 12 mm and a more challenging 6 mm beam two times each on the first test day and between two and four times on the second test day. All mice were successful in crossing the full span of each beam during all test sessions and trials (Figure 1A). Wild-type mice crossed the 12 mm beam in 2.6 to 11.3 seconds (mean = 5.06 ± 0.29 seconds), while *Vglut3-ko* mice crossed in 2.2 to 35.6 seconds (mean = 8.9 ± 0.82 seconds). Although the traversal time on the 12 mm beam was on average faster for the wild-type group, the fastest mouse from the two cohorts was a P73 *Vglut3-ko* mouse that crossed the 12 mm beam in 2.2 seconds. Traversal time on the thinner 6 mm beam was slower than the 12 mm beam in both wild-type (mean = 12.3 +/- 0.99 s and *Vglut3-ko* mice (mean = 15.3 +/- 0.88 s).

**Figure 1.**
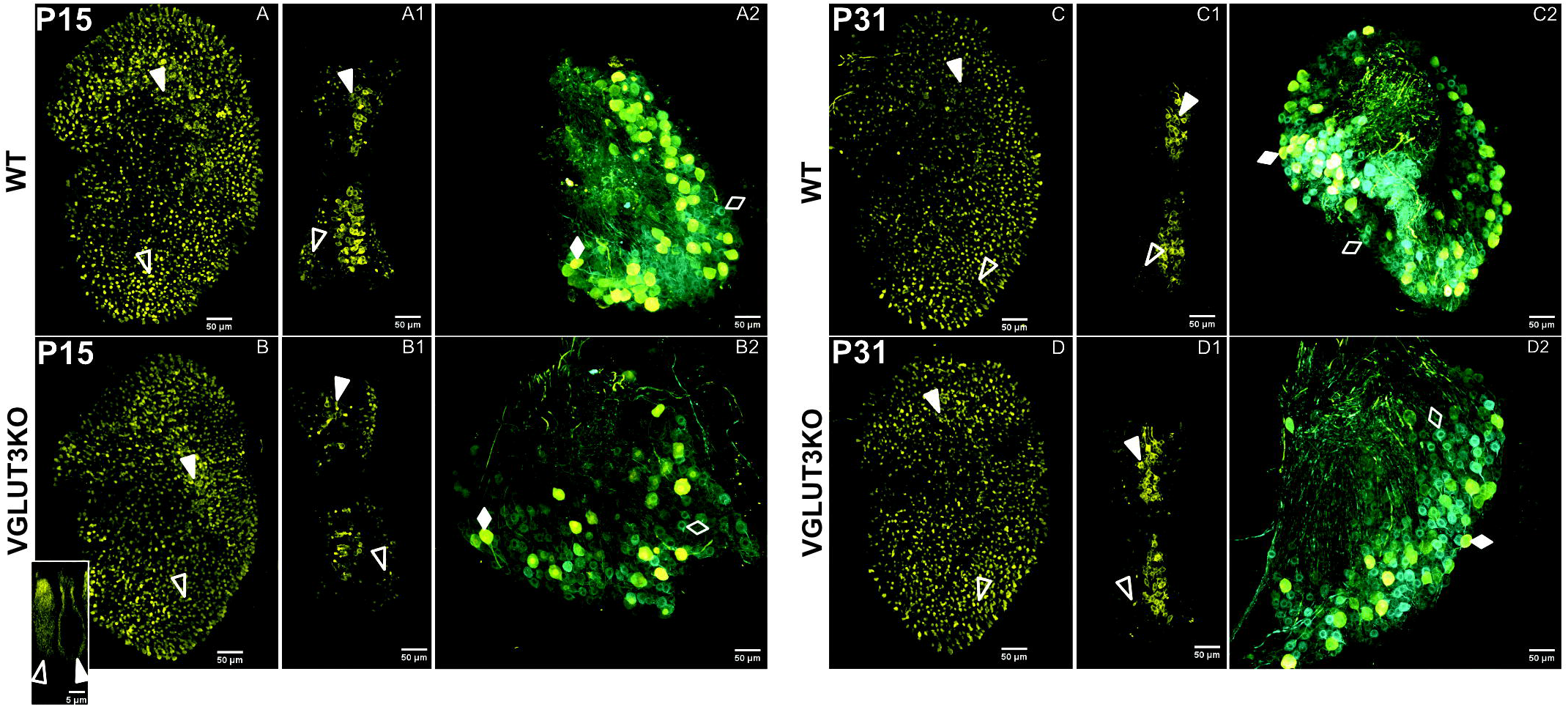
Balance function is largely intact in VGlut3KO animals though some knockout animals take longer to cross an easy beam than wild-type animals. **A.** Traversal times for a 6 mm and 12 mm beam measured in C57B6J wild-type (blue, n=11) and *Vglut3-ko* mice (red, n=15). Each mouse was tested twice each on two test days separated by one day (circles and crosses, respectively). Some animals crossed the 6 mm balance beam 4 times on the second test day. **B**. Distribution of traversal times displayed according to age, strain, and beam width. **C**. Least-means square plots of traversal time plus 95% confidence intervals by age range for wildtype and knockout mice. Traversal times are longer for the knockout mice, but this is not dependent on age. **D**. Least-means square plots of traversal time plus 95% confidence intervals by beam width for wildtype and knockout mice. Vglut3 KO mice have longer traversal times but only for the 12 mm condition. E Number of slips and stalls by strain and beam width.

We performed a repeated measures two-factor analysis of variance with two between-subject factors (age, and strain) and one within-subject factor (beam width) (Figure 1B, full distribution of data shown by grouping factors), which found that traversal time was not dependent on age (p = 0.69) but was dependent on both strain (p=0.0148) and beam width (p<0.001) (Figure 1 C&D least-means squared plus 95% confidence intervals). Both strains took longer to cross the 6 mm beam than the 12 mm beam and performance on the narrower beam was more variable between animals, strains, and across trials for an individual animal. Whereas the 6 mm beam took longer for both strains to cross, strain-dependent differences in traversal time were notable for the 12 mm beam but not the 6 mm beam, with the *Vglut3-ko* animals taking longer to cross the 12 mm beam than the wildtype animals (Figure 1D).

Part of the difference in traversal times may be related to more frequent stalls and slips in some *Vglut3-ko* mice when compared to wild-type mice (Figure 1E). This difference is most obvious on the 12 mm beam, whereas the wild-type mice exhibited no slips and few stalls. While both strains exhibited some stalls and slips on the narrower 6 mm beam some, *Vglut3-ko* mice performed several slips and stalls, even on the wider 12 mm beam. Notably, the stalling in the *Vglut3-ko* was sometimes accompanied by urination and defecation, perhaps indicating a heightened anxiety response.

### 3.2 The maturation and organization of vestibular epithelia in VGLUT3 knockout mice is similar to that in wild-type mice

We used immunolabeling to examine the regional organization of afferent fibers and hair cells in the utricle and anterior cristae of wildtype and *Vglut3-ko* mice. Analyzed tissue included immature and mature time points (P15 and P31, respectively) to assess epithelial and neural maturation in the context of the reduced input in *Vglut3-ko* mice.

#### Calretinin labels central zone calices and peripheral Type II hair cells in the *Vglut3-ko* mouse

In rodents, calretinin expression is absent at birth and emerges in the pure-calyx afferents in the central zones during the 2^nd^ post-natal week. Since the expression pattern for calretinin is altered in SGN of *Vglut3-ko* mice that lack glutamatergic input, we hypothesized that the late emergence of calretinin in vestibular afferents may also be activity driven and thus impacted by the altered inputs of the *Vglut3-ko* mice. As shown in Figure 2, patterns of calretinin expression in the vestibular epithelia and ganglion of knockout mice are normal, as in the wild-type epithelia. We present immunolabeling for calretinin in utricles, cristae, and vestibular ganglia from wild type and *Vglut3-ko* mice at P15 (Figure 2A through 2B2) and P31 (Figure 2C through 2D2). Calretinin stains neuronal afferent calices in the central zones (solid white triangles) and most, but not all, Type II hair cells in the peripheral zones (hollow white triangles). In the vestibular ganglion, calretinin labels a subset of all β-III Tubulin-positive soma in both the wild type and *Vglut3-ko* mice (figure 2A2, figure 2B2, respectively). A similar result is seen in the functionally mature *Vglut3-ko* epithelia and ganglion. Figures 2C, through 2C2 show the calretinin expression in the wild type utricle (fig. 2C), crista (fig. 2C1) and ganglion (fig. 2C2) at P31. Overall, the differential pattern for calretinin expression, a hallmark of vestibular neuron diversity, is normal in *Vglut3-ko* mice.

**Figure 2.**
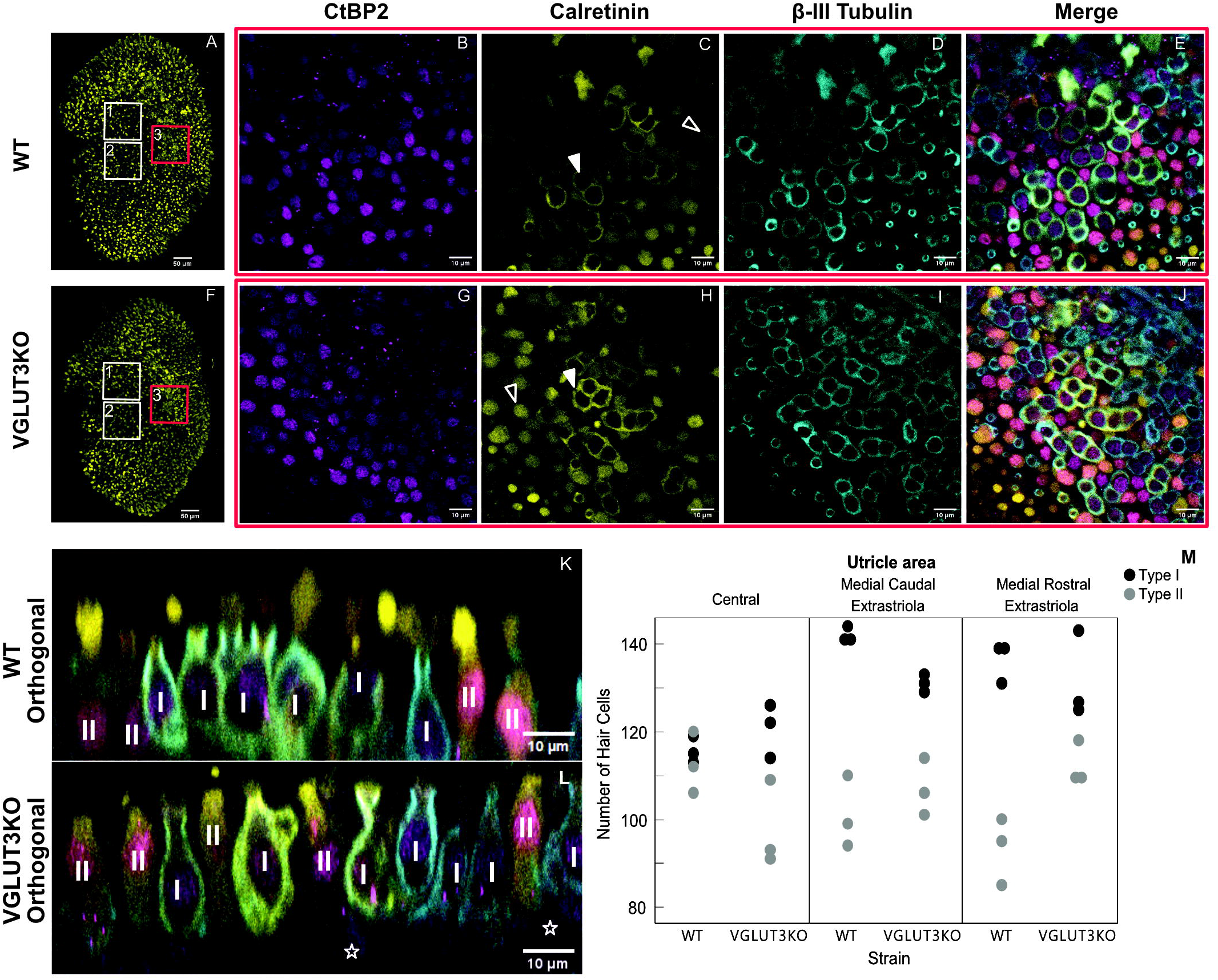
Calretinin expression in hair cells and ganglion neurons develops normally in VGlut3KO mice. Immunolabeling to visualize calretinin expression in vestibular epithelia and superior vestibular ganglia of WT and Vglut3KO mice at p15 (**A** through **B2**) and P31 (**C** through **D2**). At both ages and in both the WT and VGlut3KO mice, calretinin staining is robust in many central zone calyces of the utricle and crista and in most type II hair cells in the peripheral zones. In the ganglion, calretinin is enriched in a subset of all somata. **A** through **A2**: The pattern of calretinin expression (yellow) at P15 in WT utricle (**A**), crista (**A1**) and ganglion (**A2**). In panels **A** and **A1**, Calretinin-labels type II hair cells as well as calyces. White hollow triangles point to type II hair cells in the peripheral zones. In panel **A2**, all somata label for TUJ1 (blue), but only a subset are enriched for both TUJ1 and Calretinin. Filled white diamonds point to an example soma from neurons that co-express calretinin and TUJ1. Empty white diamonds point to an example soma from neurons that express TUJ1 but not calretinin. **B** through **B2**: The pattern of calretinin expression at P15 in Vglut3KO. Symbols same as in **A**. **Inset in B**: orthogonal view from utricular central zone of a calretinin-labeled type II hair cell (hollow white triangle) and a calretinin-labeled calyx (solid white triangle) in a VGlut3KO. **C** through **C2**: The pattern of calretinin expression at P31 in WT utricle (**C**), crista (**C1**) and ganglion (**C2**). Symbols same as in **A**. **D** through **D2**: The pattern of calretinin expression at p31 in Vglut3KO utricle (**D**), crista (**D1**), and ganglion (**D2**). The expression pattern is similar to that seen in the WT. Except for the inset, all images are maximum intensity projections at 10x with a scale bar of 50 µm. Inset is an orthogonal slice from a Vglut3KO utricle at 63x, with a scale bar of 5 µm.

### *Vglut3-ko* mice have similar numbers of Type I and Type II hair cells as wild-type mice

Previous studies have reported that SGN diversity in the *Vglut3-ko* is impacted by the lack of glutamatergic input, resulting in distorted populations of SGN Type I subgroups as identified by SGN expression of calcium-binding protein calretinin (Shrestha et al., 2018). We predicted that this lack of input in *Vglut3-ko* mice would also impact vestibular epithelia. Type II vestibular hair cells do not synapse with calyceal afferent neurons and are thought to rely solely on quantal glutamatergic transmission. We therefore considered whether the *Vglut3-ko* would have a reduced number of Type II hair cells, and a compensatory increase in Type I hair cells. To investigate, we obtained high-magnification scans of wildtype and *Vglut3-ko* mice (figure 3A – 3L).

**Figure 3.**
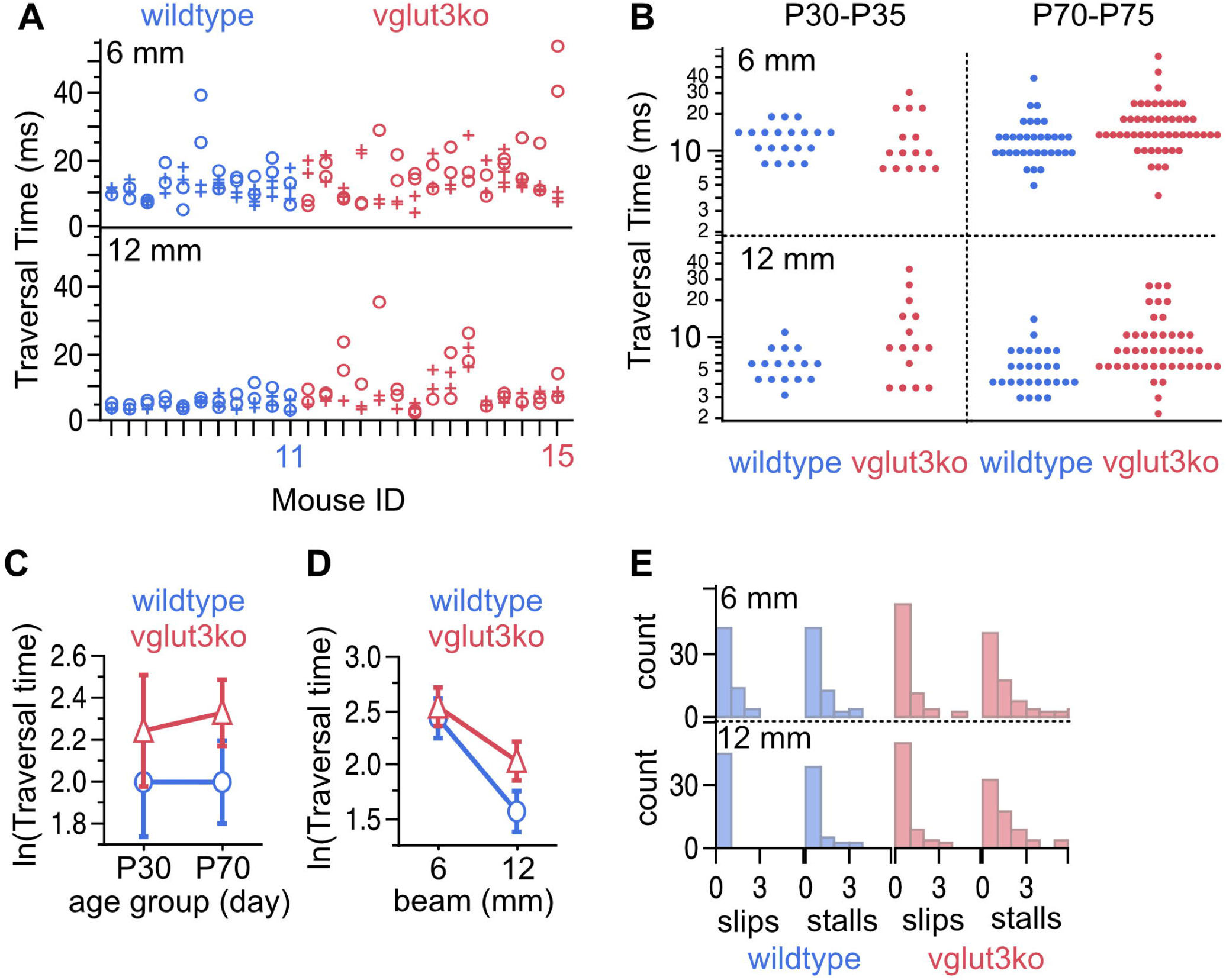
Type II hair cells in the *Vglut3-ko* are present in similar numbers as in wild-type mice. High magnification scans of wild type (WT) and *Vglut3-ko* utricle at P15 were used to quantify vestibular hair cells in three regions. **A**: Utricle from WT mouse aged P15. The boxes overlayed outline the approximate location of the high magnification scans used in the hair cell quantification: the medial rostral extrastriola (white box labeled ‘1’), medial caudal extrastriola (white box labeled ‘2’) and the central zone encompassing the striola and juxtastriola (box labeled ‘3’ and outlined in red). **B** through **E**: High magnification scan from central zone of the utricle (red box 3), immunolabeling with CtBP2 in magenta (**B**), calretinin in yellow (**C**), β-III Tubulin in blue (**D**), and the merged scan (**E**). In **C**, solid white triangles point to example calices in the striolar zone, and hollow white triangles point to example Type II hair cells in the extrastriolar zone. **F** through **J:** Same organizational layout and symbols as **A** through **E**, but for *Vglut3-ko* utricle aged P15. **K** and **L**: Orthogonal view of WT (**K**) and *Vglut3-ko* (**L**) utricle through the striolar zone. Type I and Type II hair cells are identified. Hollow white stars on **L** label example supporting cells. **M**: Quantification of Type I and Type II hair cells at P15 in the medial rostral extrastriola, medial caudal extrastriola, and the central zone encompassing the striola and juxtastriola (n= 3 utricles per strain, 6 animals total). Each scan area was approximately 1.03 x 10^-2^ mm^2^. All images except for A and F are single slices at 63x with a scale bar of 10 µm. **A** and **F** are maximum intensity projections at 10x with a scale bar of 50 µm.

Confocal scans taken at P15 through the utricles of immunolabeled wild-type (figure 3A -E) and *Vglut3-ko* mice (Figure 3 F-L) show no difference in the organization of the epithelia. Figure 3A and 3F show low-magnification scans of utricles labeled with calretinin, which typically labels Type II hair cells and the pure calyx-terminals of central zone afferents. White boxes on Figures 3A and 3F outline the approximate areas of utricles that were scanned and quantified, with box 1 outlining the medial rostral extrastriola, box 2 outlining the medial caudal extrastriola, and box 3 outlining the central zone which includes the striola and juxtastriolar regions. Figures 3B-3E (wild type) and 3G-3J (knockout) show representative high-magnification scans through the central zone with antibodies against CtBP2 to label nuclei and synaptic ribbons, antibodies against calretinin to label most Type II hair cells and post-synaptic calyces in the striola, and antibodies against β-III Tubulin cup to label all neurons. Synaptic ribbons were ample in all types of hair cells and in all three regions of the utricle in wild-type (Figure 3B) and knockout (Figure 3G) animals. We did not quantify ribbons, because we found that ribbons in the *Vglut3-ko* samples appeared clumpy and in very tight clusters making it difficult to distinguish between individual ribbons. This is consistent with the observation in *Vglut3-ko*’s inner hair cells where electron microscopy found abnormally shaped ribbons (Seal et al., 2008). Whether there is a similar distortion in ribbon morphology in vestibular hair cells remains to be determined. Calretinin expression in both the wild type (Figure 3C) and the *Vglut3-ko* (Figure 3H) was robust and labeled calices in the striola (white triangles), as well as Type II hair cells outside of the striola (hollow triangles).

β-III Tubulin was found in all calyceal afferents and was thus used for counting Type I hair cells in wildtype (Figure 3D) and *Vglut3-ko* (Figure 3I) scans. CtBP2 nuclei at the hair cell level were counted as type II hair cells when not surrounded by a β-III Tubulin cup. Note examples of Type I and Type II hair cells in high-magnification orthogonal views (Figures 3K and 3L from a wild-type and *Vglut3-ko* utricle, respectively). Supporting cell nuclei are found deeper in the tissue and are thus easily distinguished from hair cell nuclei in orthogonal views of the tissue (hollow stars Figure 3L). Based on these criteria we counted the number of Type I and Type II hair cells by epithelial region in in three biological replicate utricles in *Vglut3-ko* and wild type. For more details, see Methods.

We found that knocking out VGLUT3 did not dramatically alter the number of Type I and Type II hair cells in the utricle, although minor regional differences were observed (Figure 3M, see also table 1 for averages). As was the case in the wild type, the *Vglut3-ko* had a greater number of Type I hair cells (Figure 3M, black dots) than Type II hair cells (Figure 3M, gray dots) in all three regions analyzed. The lack of quantal input did not lead to a decrease in the number of Type II hair cells in the *Vglut3-ko*. Indeed, the number of Type II hair cells in the extrastriolar regions was slightly higher in the knockout than in the wild type, which was contrary to our expectations. We therefore believe the slight differences in numbers of Type I and Type II hair cells across the regions reflect natural variations between the strains.

**Table 1.**

| Strain | n | Measure | Utricle Area |  |  |
| --- | --- | --- | --- | --- | --- |
|  |  |  | Central | Medial Caudal<br>Extrastriola | Medial Rostral<br>Extrastriola |
| C57BL6 | 3 | Ratio Type I:Type II | 1.03 ± 0.04 | 1.41 ± 0.06 | 1.46 ± 0.04 |
|  | 3 | Type I count | 115.67 ± 1.76 | 142 ± 1.00 | 136.33 ± 2.67 |
|  | 3 | Type II count | 112.67 ± 4.06 | 101.00 ± 4.73 | 93.33 ± 4.41 |
|  | 3 | Total Number of Hair<br>Cells | 228.33 ± 4.81 | 243.00 ± 5.69 | 229.67 ± 6.98 |
|  | 3 | Density per 100 µm<sup>2</sup> | 2.22 ± 0.05 | 2.36 ± 0.06 | 2.23 ± 0.07 |
| Vglut3-ko | 3 | Ratio Type I:Type II | 1.24 ± 0.07 | 1.23 ± 0.05 | 1.17 ± 0.07 |
|  | 3 | Type I count | 120.67 ± 3.53 | 131.00 ± 1.15 | 131.33 ± 5.84 |
|  | 3 | Type II count | 97.67 ± 5.70 | 107.00 ± 3.79 | 112.33 ± 2.85 |
|  | 3 | Total Number of Hair<br>Cells | 218.33 ± 7.51 | 238.00 ± 2.65 | 243.67 ± 5.49 |
|  | 3 | Density per 100 µm<sup>2</sup> | 2.12 ± 0.07 | 2.31 ± 0.03 | 2.37 ± 0.05 |

Values are means ± standard error Results collected from scans 1.03 x 10<sup>-2</sup> mm<sup>2</sup>

### Enriched expression for KCNQ4 in the central zone calyces is normal in the knockout mouse

KCNQ4, a subunit of low-voltage gated Kv7 channels is known to be localized to afferent terminals in the central zones of vestibular epithelia (Spitzmaul et al., 2013). Much like calretinin expression, KCNQ4 channels and Kv7-like currents emerge in the central zones over the first two weeks of development (Rocha-Sanchez et al., 2007; Meredith and Rennie, 2015). Figure 4 shows maximum-intensity projections of confocal scans through a utricle at P15 and semicircular canal at P31 in a *Vglut3-ko* and wild-type mice. The boundaries of the epithelial region are demarcated by a dashed line. As evident in both the knockout and wildtype samples, KCNQ4 is enriched in the calyces (cup-like labeling) in the central zones of utricle and cristae samples but not in the peripheral zones. Thus, the lack of glutamatergic input does not alter the developmental emergence of KCNQ4 subunits.

**Figure 4.**
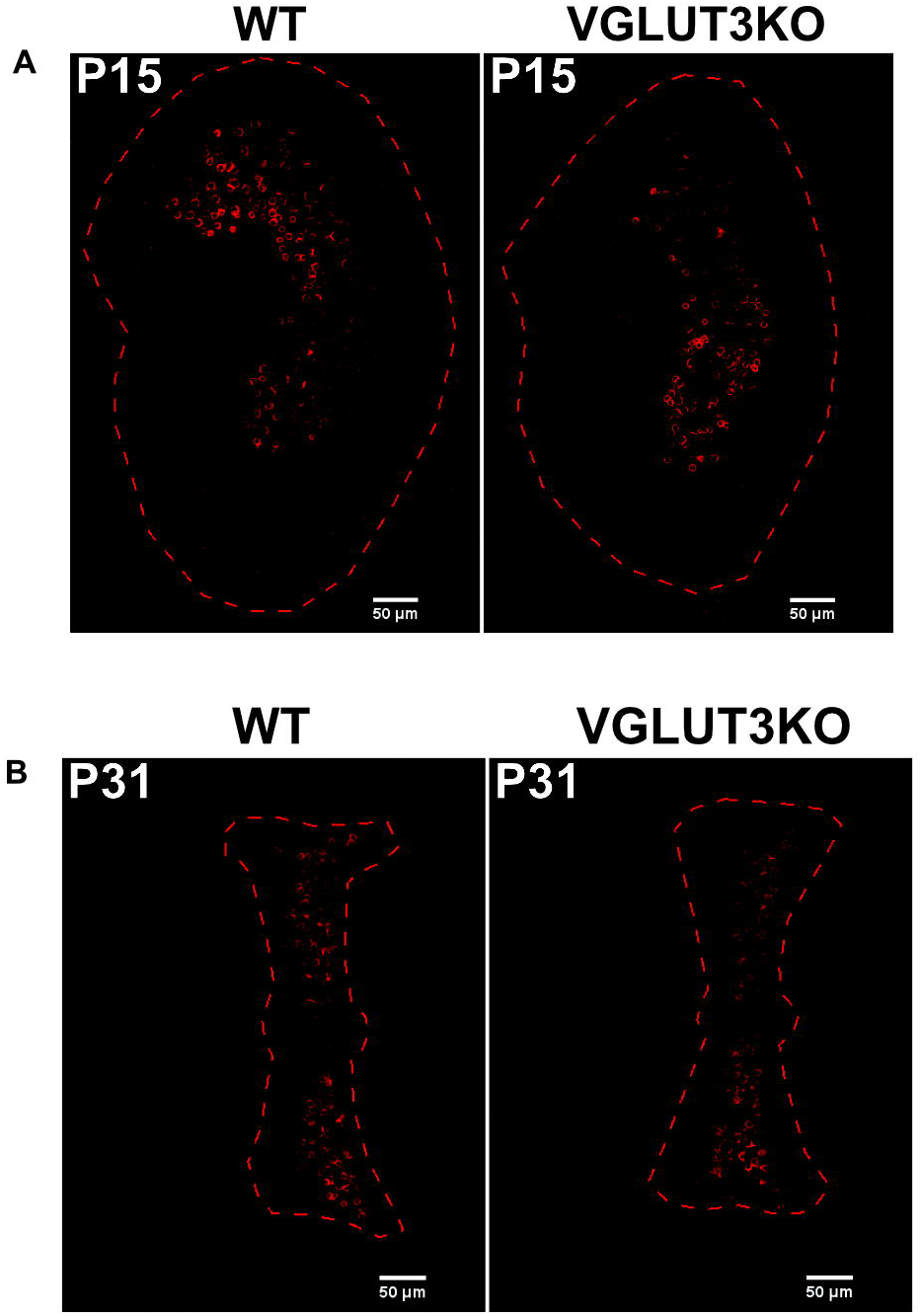
KCNQ4 expression in vestibular epithelia is consistent with WT mice. **A,B**: Immunolabeling for KCNQ4 in example utricles at P15 (**A**) and canal cristae at P31 (**B**) in WT and *Vglut3-ko*.

### 3.3 *Vglut3-ko* mice have normal biophysical diversity

Although immunohistochemistry suggests that KCNQ4 emerges normally in the central zones of the vestibular epithelia, we considered whether expression levels of these and other channels were in fact normal. We did so by comparing the firing patterns and whole-cell currents of vestibular neurons in *Vglut3-ko* and wild-type animals. Given that SGN in *Vglut3-ko* mice are hyper-excitable (Babola et al., 2018) we hypothesized that vestibular afferent somata would similarly exhibit an increased level of excitability relative to wildtype mice. Below, we present whole-cell, perforated-patch clamp recordings from cultured dissociated somata of vestibular afferent neurons from *Vglut3-ko* mice aged post-natal day (P)8 to P23 (n=70) and wild-type mice of similar ages (P9-P24, n=71).

#### *Vglut3-ko* mice retain firing pattern diversity

Figure 5 presents the range of firing patterns found in *Vglut3-ko* VGN (Figures 5A through 5E). Spikes were generated in response to 500 millisecond square pulse current injections at different amplitudes. Below each trace is a plot depicting the spike count as a function of current level (see Methods for details). The diversity of firing patterns is similar to that previously described in in wildtype mouse and rat leading us to classify the patterns according to the scheme previously described (Kalluri et al., 2010; Ventura and Kalluri, 2019; Kalluri, 2021; Bronson and Kalluri, 2023). Transient-firing VGN produced a single action potential in response to current injections that either reach (black trace) or exceed (grey) spike threshold (Figure 5A1). Note that the level dependence of the example transient-firing VGN was such that a single action potential was generated across all current steps (Figure 5A2, black and grey circles correspond to traces shown in Figure 5A1). A handful of VGN failed to generate an action potential in response to any current step and were classified as graded-response VGN (Figure 5B1). A wide-range of firing patterns were broadly classified as sub-groups of sustained-firing. Sustained-A-firing neurons fired continuously throughout the current step near threshold (Figure 5C1, black traces). The level dependence of sustained-A-firing VGN varied such the same suprathreshold current injection generated more spikes in some sustained-firing VGN than in others (left, grey trace versus right, grey trace). Compare left and right plots in figure 5C2 for the level dependence of the spike patterns in the two example sustained-A *Vglut3-ko* VGN shown in 5C1. Sustained-B neurons fired at least two action potentials that often increase in number at larger current steps (Figures 5D1 and 5D2). Sustained-C neurons fired a single action potential followed by voltage oscillations (Figure 5E1).

**Figure 5.**
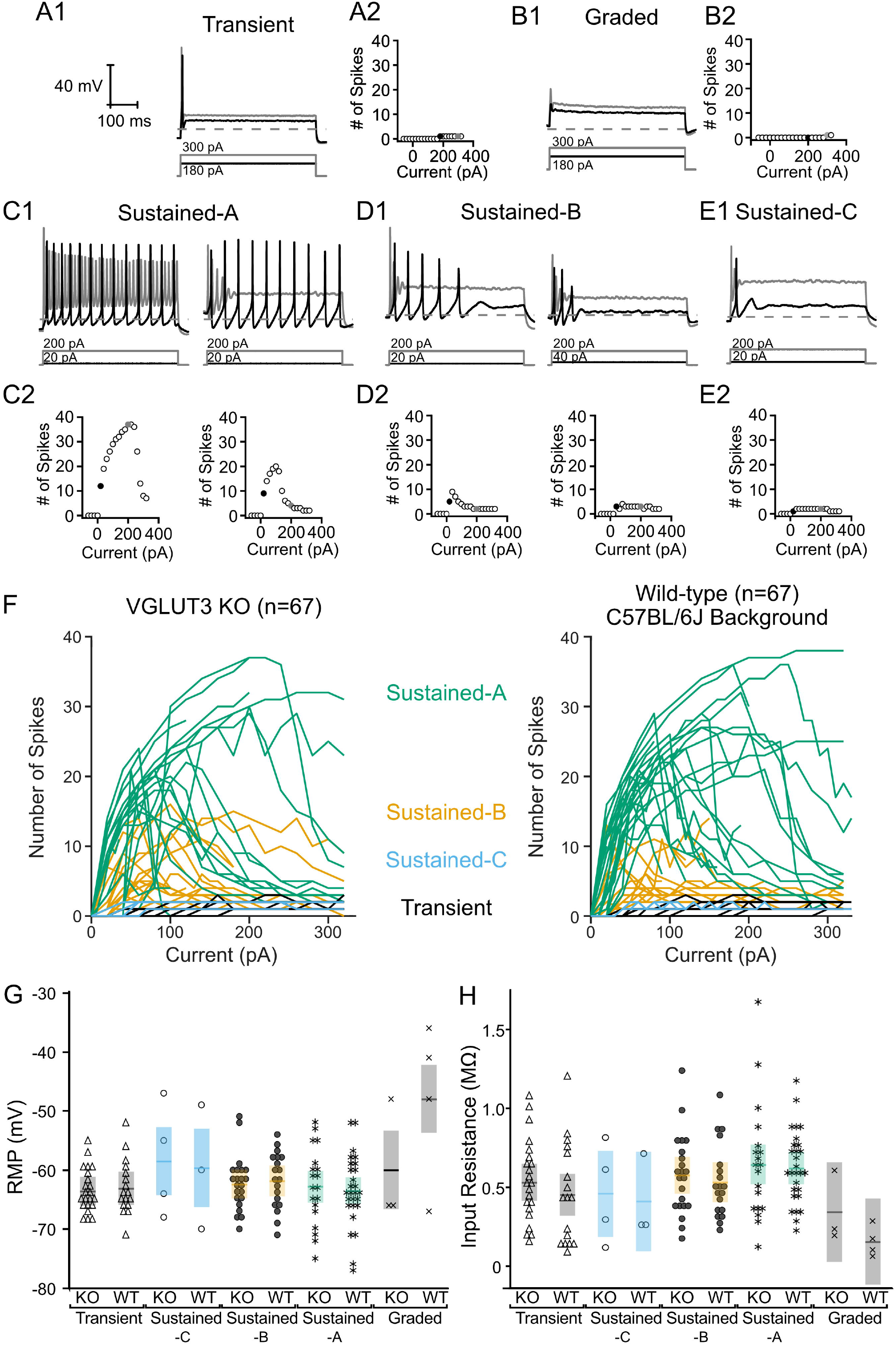
*Vglut3-ko* mice retain heterogenous depolarization-evoked firing patterns found in control mice. All cells shown extracted from *Vglut3-ko* mice. Scale bars are consistent throughout all traces. Depolarizing current steps shown in black are the closest 20 pA current step to threshold in each trace, response to superthreshold 200-300 pA current steps shown in gray. **(A1)** Transient-firing neurons fire a single action potential at the onset of suprathreshold depolarizing current steps. Dashed line indicates -60 mV. The amplitude of the positive current step necessary to reach threshold is shown below each trace in black and larger current step shown in grey. **(A2)** Relationship between size of current injection and number of spikes generated with black and grey traces indicated by black and grey filled circles, respectively **(B1)** Graded cells did not fire action potentials in response to depolarizing current steps regardless of current amplitude. **(B2).** Automated spike detection algorithm occasionally detected spikes in response to large current steps, but these were ignored as all spikes were verified manually. **(C1).** Sustained-A neurons fire continuously throughout the current step. However, neurons classified as sustained-A often varied in the number of spikes generated across in response to larger current injections. Left: sustained-A neuron that fired continuously at threshold in black with no accommodation in response to larger current injections in grey. Right: Cell fired that continuously at threshold (black) and accommodated in response to larger current injections (gray). **(C2)** Range of current responses show similar number of spikes generated at threshold (black filled circle) with significantly fewer spikes generated in response to 200 pA depolarizing current injection. **(D1)** Sustained-B neurons fire multiple action potentials after the onset of the depolarizing current steps and are rapidly-adapting. Left: Highly excitable sustained-B neuron. Right: Less excitable neuron with higher threshold current. **(D2)** Number of spikes generated in response to range of current injections in neurons shown in above traces. **(E1).** Sustained-C neurons fire a single action potential followed by voltage oscillations. **(E2).** Spike detection algorithm occasionally detected voltage oscillations as spikes. (**F)** Number of spikes generated by *Vglut3-ko* (left, n=67) and wild-type cells (right, n= 67). Each line represents a single cell. Lines are color-coded according to their firing pattern with sustained-A in green, sustained-B in orange, sustained-C in blue and transient cells in black. **(G)** There was no significant difference in resting membrane potential (RMP) and **(H)** input resistance between *Vglut3-ko* and wild-type mice.

To compare the firing patterns in *Vglut3-ko* and wild-type VGN, Figure 5F shows the level dependence of spike number in 67 *Vglut3-ko* VGN (left) and 67 wild-type VGN (right). As current injections increase, sustained-A firing VGN (green) in *Vglut3-ko* and wild-type VGN show the same maximum number of spikes, similar patterns of increasing spike number followed by accommodation. Similarly, sustained-B firing VGN (orange) in both *Vglut3-ko* and wild types generate the same maximum number of spikes and accommodate at roughly the same current levels. The sustained-C firing (blue) and transient firing VGN (black) do not show level dependence because they generate a single action potential, but cells from *Vglut3-ko* and wild-type reach threshold at similar current levels. These results demonstrate that VGN excitability as measured by the level dependence of spike number is similar in both *Vglut3-ko* and wild-type mice.

Resting membrane potential (RMP) and input resistance were not different between VGN in *Vglut3-ko* and wild-type mice. Figure 5G shows the resting membrane potential in VGN from both *Vglut3-ko* (grey) and wild-type mice (black). A two-factor analysis of variance with two between-subject factors (firing pattern and strain) showed that RMP was not dependent on strain (p = 0.1146) but was dependent on firing pattern (p<.0001). In both *Vglut3-ko* and wild-type mice, RMPs in graded-response VGN were more depolarized than those in sustained-A (p<.0001), sustained-B (p=0.003), and transient-firing VGN (p<.0001). However, we did not find any difference in RMP between VGN of *Vglut3-ko* and wild-type mice overall or between VGN of any firing pattern. As was the case with RMP, input resistance was dependent on firing pattern (p=0.018), but not strain (p=0.25). Figure 5H shows the input resistance in *Vglut3-ko* (grey) and wild-type mice (black). In both *Vglut3-ko* and wild-type mice, graded-response VGN exhibited lower input resistance and excitability than sustained-A VGN (p=0.0161). We found that input resistance in VGN of *Vglut3-ko* and wild-type mice were equal across all firing patterns. Overall, these results show that the resting conductances in VGN of *Vglut3-ko* and wild-type mice are not different.

### *Vglut3-ko* mice show no obvious differences in VGN firing pattern and cell size between first through third week of development

Having shown that all VGN firing patterns are found in the *Vglut3-ko* mice and have similar properties, we next examined the proportion of VGN that fall within these categories through development. Figures 6A and 6B show the proportion of neurons classified into each firing pattern category at three age groups: before P11 (N_WT_=16, N_KO_=19), between P12-P18 (N_WT_=36, N_KO_=29) and after P19 (N_WT_=19, N_KO_=22) in wild type and *Vglut3-ko*, respectively. Figures 6C and 6D show the same data but with the exact ages and firing pattern for each cell that contributes to the data in 6A and 6B (the boundaries for the age bins are indicated by the dashed lines). As the vestibular system matures in wildtype mice, transient-firing VGN became more prevalent (22% to 32%) while sustained-A firing neurons became less prevalent (47% to 16%). We also observed a developmental increase in the number of Sustained-C firing and graded responses, both were not observed at the earliest time point in mouse but have been observed at early time points in a previous study in rat).

**Figure 6.**
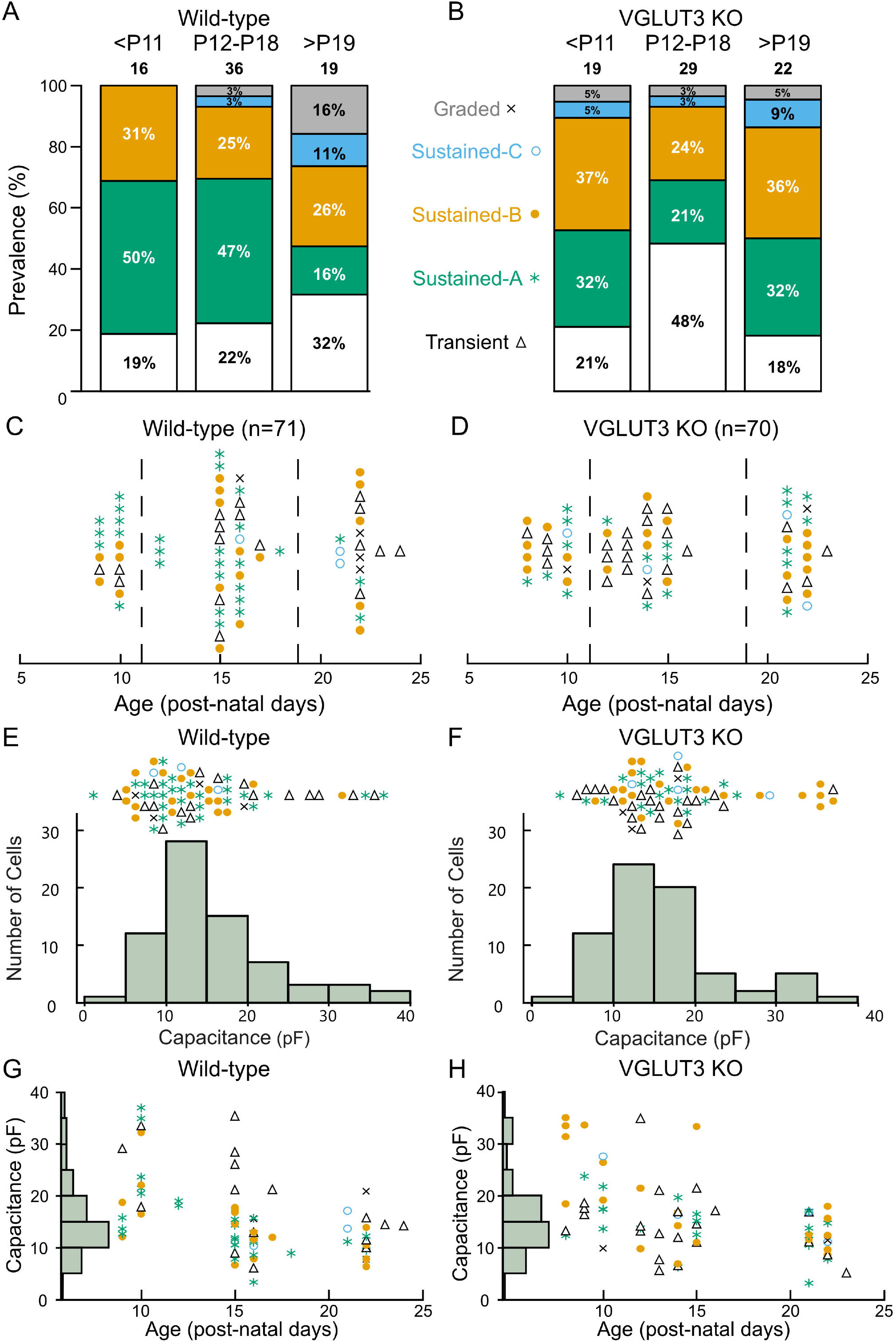
Distribution of firing pattern and cell size in*Vglut3-ko* mice is consistent with wild-type. Cells were recorded between according to days post-natal (P) 8 to 24. **(A)** Proportion of cells of each firing pattern at three periods of early development: P8-10 (<P11), P12-18 and P21- 24 (>P19) in the wild type. **(B)** Proportion of cells of each firing pattern in the *Vglut3-ko*. Note that cells of all firing patterns are seen at each period of development in both *Vglut3-ko* and wild-type mice except graded and sustained-C, which are less numerous overall **(C)** Every *Vglut3-ko* cell (N=71) with transient-firing patterns (black triangles), sustained-A firing patterns (green asterisks), sustained-B firing patterns (orange filled circles), sustained-C firing patterns (open blue circles) and graded firing patterns (indicated by X) organized by age in post-natal days. Dashed lines indicate age boundaries used in figures 6A and 6B. **(D)** Wild- type cells (N=70) and their firing pattern organized by age. **(E)** Size distribution of all *Vglut3-ko* neurons was measured from membrane capacitance (*C_m_*) and shown below. Each cell and corresponding firing pattern is arranged according to its size above. **(F)** Size distribution (below) and firing pattern (above) of wild-type VGN. **(G)** *Vglut3-ko* cell size is shown as a function of age. Note the absence of large cells recorded after P15. **(H)** Wild-type cell size plotted as a function of age. Large cells are similarly absent in recording of wild-type neurons.

The age-dependent increase in transient-firing patterns and decline in sustained-A firing patterns is reminiscent of a similar change in firing pattern prevalence that we’ve previously reported in rat and attributed to a developmental up-regulation in low-voltage gated potassium currents (Kalluri et al., 2010; Ventura and Kalluri, 2019; Bronson and Kalluri, 2023). Note however that the shift towards the transient-firing pattern is more dramatic in rat where only about 5% of VGN in the third post-natal week have sustained-A firing patterns (Ventura and Kalluri, 2019). Together, these results suggest that VGN in the wild-type mouse gradually decrease in excitability as the vestibular system matures, possibly due to the upregulation of low-voltage gated potassium conductances such as those mediated by KCNQ.

We found all major firing pattern groups in each of the age groups in the *Vglut3-ko* animals (Figure 6B & D). Like the developmental trend in wild-type mouse and rat VGN, we saw an increase in the prevalence of transient-firing patterns between the first and second post-natal weeks (21% to 48%). However, by the third to fourth post-natal weeks, nearly 32% of the population in had a sustained-A firing pattern (as compared to the 16% in wild-type mouse and 5% in rat). At first glance, this increase in the prevalence of sustained-A firing could mean that the lack glutamatergic input has pushed some VGN towards increased excitability, as previously reported in the SGN (Shrestha et al., 2018). However, this interpretation is complicated by technical issues that created a sampling bias in our dataset toward small soma in the third/fourth post-natal weeks (as discussed below).

We previously showed in rat that VGN cell size is bimodally distributed with a main mode of small-to medium sized soma comprised of transient-firing and sustained-firing VGN (putative cell bodies of peripheral zone afferents) and a second mode of large soma with predominantly transient-firing patterns (putative cell bodies of central-zone irregular-firing afferents) (Kalluri et al., 2010; Bronson and Kalluri, 2023). Based on the correlation between soma size and calretinin expression (Kevetter and Leonard, 2002), large soma with transient-firing patterns likely belong to the irregularly-firing afferents that project to the central zone. As illustrated by the histogram of membrane capacitance (*C*_m_, an electrophysiological measure of somatic surface area; Limón et al., 2005), VGN soma size in both wild-type mouse and *Vglut3-ko* is bimodally distributed with a group of small cells and a handful of relatively large cells (Figure 6E & F, respectively). Since the size distribution of VGN from the *Vglut3-ko* mouse is bimodal with a similar range of sizes as the wild type VGN, the lack of glutamatergic input did not impact cell size, and we use cell size as a rough indicator of zonal identity.

Note, however, that in both wild-type and *Vglut3-ko* mice, we had very few recordings from large soma after P20, probably due to selection bias induced by technical challenges such as greater myelination and fragility of the cells (Figures 6G and 6H). Thus, while the culture plates continued to have large-sized cells in both the wild-type and *Vglut3-ko* mice, our data likely under-represents the number of large cells. This could in part account for why there are more sustained-firing VGN at the older ages. However, this technical issue was equally present in both the wild-type and *Vglut3-ko* mouse suggesting that there may still be subtle differences in the ion channel properties of specific sub-populations of neurons.

To look at this question with further granularity, we focus on the ten largest cells in our population in both the wild-type and *Vglut3-ko*. In wild-type animals, nearly half of the cells have a transient-firing pattern whereas in the *Vglut3-ko* group the firing patterns are mostly sustained-B (6/10, orange circles) or sustained-A (2/10, green asterisks). These results suggest the possibility that central zone contacting VGN in the *Vglut3-ko* remain more excitable in the absence of glutamate input.

### Large, potentially central zone contacting VGN in *Vglut3-ko* mice may have less *I*_KL_ than similarly-sized VGN in the wild type

Although KCNQ4 channel expression emerges successfully in the central zones of vestibular epithelia in *Vglut3-ko* mice, immunohistochemistry is not a quantitative measure of channel function (Figure 4). Channel densities may differ in the knockout animal as suggested by the greater prevalence of sustained-firing neurons amongst the large somata (recall Figure 6). To further interrogate this we compared the whole-cell currents generated in response to depolarizing voltage steps in a subset of 49 VGN from *Vglut3-ko* mice and 46 from the wild type that had voltage clamp recordings. Figure 7A and 7B show the size distribution and firing patterns of this subset of VGN from the *Vglut3-ko* (red, left) and wild-type (blue, right) mouse, respectively. We selected eight large cells in the *Vglut3-ko* and seven large cells in the wild type mouse (see right-pointing arrow). We also compared the whole-cell currents of small cells. Small cells were taken from a cluster of cells between 12 and 17 pF (n=21 and n=18, *Vglut3-ko* and wild type, respectively).

**Figure 7.**
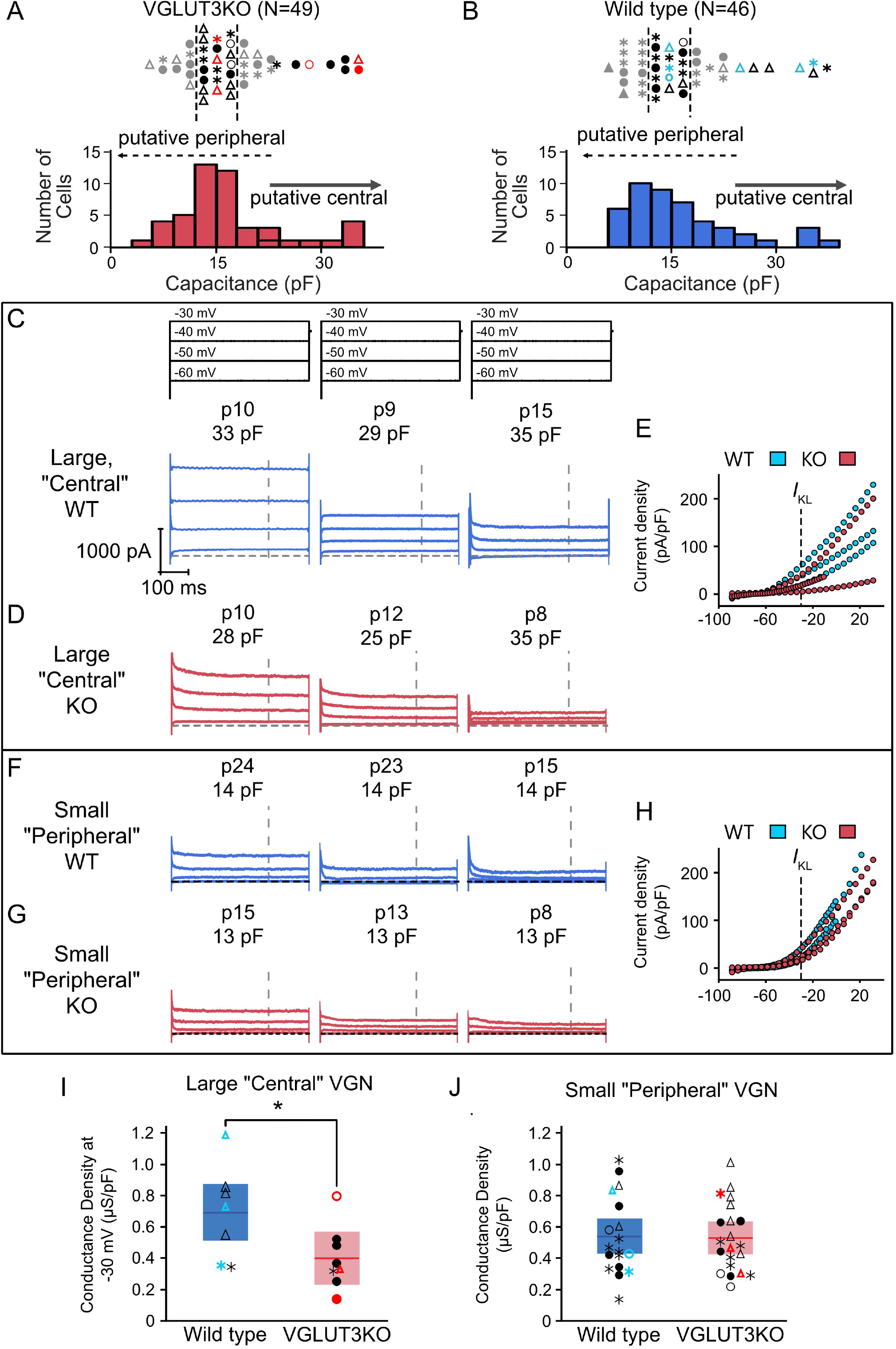
Large putative central zone contacting VGN may have reduced *I*_KL_ in *Vglut3-ko* mice. Whole-cell currents were recorded in voltage-clamp in a subset of 49 *Vglut3-ko* cells and 46 wild-type cells. **(A)** Size distribution (below) and firing pattern (above). The 8 largest cells were classified as putative central zone contacting VGN and 21 smaller cells in the largest cluster were selected as putative peripheral zone contacting VGN. Cells in red are shown in figure 7D and 7G. Grey cells were excluded. **(B)** Size distribution (below) and firing pattern (above) in wild-type VGN. 18 cells fell within the 12-17 pF size range used to classify putative peripheral contacting VGN in the *Vglut3-ko* and were classified as small VGN in the wild type. Cells in blue are shown in Figures 7C and 7F. **(C)** Cells were held at -120 mV for 30 milliseconds before stepping to -60 to -30 mV for 500 milliseconds. Cells were grouped according to their relative conductance (high, medium and low) within each size group (large putatively central VGN and small putatively peripheral VGN). Left three panels show three large wild-type VGN with currents activated from voltage-steps as above. Age and cell size as measured by membrane capacitance (*C_m_*) shown above in each trace. Dashed line indicates steady-state duration approximately 450 milliseconds after step onset during which conductance at -30 mV step is measured. **(D)** Currents, ages, cell size and current-voltage relationships of three large, putatively central VGN in *Vglut3-ko* mice. **(E)** Size-normalized current density plot across all voltages tested in the large wild-type (blue) and *Vglut3-ko* (red) VGN in figure 7C and 7D. Dotted line indicates -30 mV step where *I*_KL_-related conductance is measured. **(F)** Currents, ages, and cell sizes of three small, putatively peripheral VGN in wild-type mice. **(G)** Currents, ages, and cell size of three small, putatively peripheral VGN in *Vglut3-ko* mice. **(H)** Size-normalized current density plots across all voltages tested in the large wild-type (blue) and *Vglut3-ko* (red) VGN in figure 7F and 7G. **(I)** *I*_KL_-related conductance density at -30 mV in large VGN of both wild-type (blue) and *Vglut3-ko* (red) mice. Cells in color indicate cells shown in C-G. * p =0.0443 **(J)** Conductance density of small VGN in *Vglut3-ko* and wild-type mice.

Figures 7C through 7G shows example whole-cell current responses to depolarizing voltage steps (stimulus shown at the top of figure 7C). Figures 7C and 7D shows whole-cell currents measured in three large VGN from wild-type and three large VGN from *Vglut3-ko* mice, respectively. The cells we selected to display here were example cells with high, medium, and low net conductance density as measured at -30 mV. Current density was measured at approximately 450 milliseconds after the onset of the voltage step when the current’s reached approximate steady state (grey vertical dashed lines in figure 7C and 7D). Size normalized current densities over a range of voltages are shown to the right in figure 7E.

Figures 7F through 7H show the whole-cell currents and corresponding current densities plotted against voltage for three example small soma in the wildtype and *Vglut3-ko* mice. In contrast to the large VGN, whole cell currents are similar in both wild-type and *Vglut3-ko*.

As an estimate of low-voltage gated conductance (*I*_KL_), we computed the conductance density by dividing the current density at -30 mV by the driving force for potassium (*V*m-E_K_) in large (Figure 7I) and small VGN (Figure 7J). Large wild-type VGN have greater conductance density at -30 mV than VGN of comparable size in the *Vglut3-ko* (p=0.0443). Note that this difference is not driven entirely by the increased prevalence of large transient-firing VGN in the wild type as the transient-firing VGN in the *Vglut3-ko* had a smaller conductance density than all the transient-firing VGN in the wild-type. In contrast, Figure 7G shows that conductance density in the small VGN was equal in the *Vglut3-ko* and wild type (p=0.7922). The presence of increased *I*_KL_ in large, wild-type VGN is consistent with the finding that that the most irregular afferents project to the central zone (Goldberg, Lysakowski and Fernández, 1990). These results suggest the possibility that central projecting VGN may be more excitable in the knockout to compensate for the loss of glutamatergic input. Given the limited age range of large cells recorded in our sample, we need to gather additional data to determine whether these differences between the wild type and *Vglut3-ko* persist or perhaps increase as the vestibular system matures.

## 4. Discussion

### Glutamate release is not necessary for the diversification of vestibular ganglion neurons

Neural diversity is shaped by a combination of genetic hardwiring and activity dependent mechanisms that modify the intrinsic excitability of neurons via regulating ion channel densities during functional development (Moody and Bosma, 2005). Recent molecular analyses of spiral ganglion neurons found that, in the absence of glutamatergic input, neuronal development is stunted with neurons failing to mature and diversify into distinct molecular sub-groups (Shrestha et al., 2018; Sun et al., 2018; Sun et al., 2021). Failure to diversify is marked by hyperexcitability in response to current injections, a reduction in the RNA messenger for potassium channels, and a lack of differential expression for calretinin, a molecular marker for cell type in both the auditory and vestibular ganglion. Unlike in spiral ganglion neurons, here we found that the biophysical and molecular diversity of vestibular ganglion neurons persists even in *Vglut3-ko* mice where these neurons develop without glutamatergic input. Indicative of normal diversity, vestibular ganglion neurons in the knockout mice have a broad range of somatic sizes and have both transient- and sustained-firing patterns, with similar prevalence over maturation and similar response features as vestibular ganglion neurons in wild-type mice.

Despite lacking glutamatergic input from hair cells, *Vglut3-ko* mice were easily able to cross two different balance beams. We chose balance beam tests because they have been effective in other mouse models at detecting deficits that originate from the central/striolar zones of vestibular epithelia (Ono et al., 2020). Although some individuals within the *Vglut3-ko* cohort required a little coaxing to get started, all of them successfully crossed the balance beams. Their hesitance was likely an anxiety response to the novel experience of the beam instead of a vestibular deficit since *Vglut3-ko* mice are also known to be more anxious than wild-type mice (Amilhon et al., 2010; Horváth et al., 2018). It has been shown previously that after an acclimation period to reduce novelty in open field tests, *Vglut3-ko* mice display activity similar to wild-type controls (Horváth et al., 2018). Consistent with the idea that the delayed crossing is not a result of sensory deficit, the fastest mouse in the testing cohort was from the *Vglut3-ko* group.

Calretinin and KCNQ4 channel expression is normal and appropriately localized to afferent terminals in the central zones of vestibular epithelia in the *Vglut3-ko* mice, as previously described in wildtype rodents (Kharkovets et al., 2000; Leonard and Kevetter, 2002; Desai et al., 2005; Rocha-Sanchez et al., 2007; Spitzmaul et al., 2013). These patterns of expression emerge appropriately during development, are present by P15, and persist into maturity. This means that the lack of glutamatergic input in the *Vglut3-ko* mouse does not alter or delay key milestones of epithelial maturation. While *Vglut3-ko* animals are deaf, they did not display obvious balance dysfunctions. Thus, unlike in spiral ganglion neurons, the biophysical and molecular diversification of vestibular ganglion neurons, and the development of vestibular function (to the extent we tested it), is not critically dependent on glutamatergic input from hair cells.

Since all vestibular hair cells have many glutamatergic ribbon synapses each, it is remarkable to consider that vestibular function and epithelial organization is relatively normal in the Vglut3 KO mice. The lack of obvious vestibular behavioral deficits in the absence of glutamatergic input is consistent with previous findings in Cav1.3 and Otoferlin knockout mice, both of which have diminished synaptic function but relatively minor behavioral deficits (Platzer et al., 2000; Brandt et al., 2003; Roux et al., 2006; Dulon et al., 2009). Though vestibular deficit is not always easily detected in common vestibular behavioral tests, recent work has shown that vestibular compound action potentials are largely intact even when AMPA receptors are pharmacologically blocked (Pastras et al., 2023), indicating that the synchronous response of the vestibular nerve is not critically reliant on glutamate release. Taken together, the observations from these different mouse models and manipulations reinforce the idea that vestibular hair cells communicate by more than glutamate release (as discussed further below).

### Non-quantal transmission may provide sufficient hair cell input to drive maturation and diversification

Two types of hair cells are found throughout the vestibular epithelium. Type I hair cells are capable of transmitting by both quantal (Bonsacquet et al., 2006; Rennie and Streeter, 2006; Holt et al., 2007; Songer and Eatock, 2013; Sadeghi et al., 2014) and nonquantal mechanisms (Yamashita et al., 1990; Holt et al., 2007; Songer and Eatock, 2013; Contini et al., 2017) whereas Type II hair cells are likely to communicate by conventional quantal transmission alone. The calyx terminal is specialized to encode transient and fast head movements, and the idea that these specializations include non-conventional/non-quantal mechanisms has long been entertained in the vestibular literature (e.g., Yamashita et al., 1990). Compared to Type II hair cells, Type I hair cells have much smaller readily releasable and sustained vesicle pools (Spaiardi et al., 2022), which suggests that they may be non-quantal specialists with less reliance on quantal transmission. Non-quantal transmission as recently described relies on the unique environment created by the tight cleft between a Type I hair cell and post-synaptic calyx terminals, and the presence of special low-voltage activated ion channels in the cleft facing membranes of the hair cell and calyx terminal (Songer and Eatock, 2013; Contini et al., 2017). The unique environment is thought to create a combination of potassium accumulation in the cleft, tonic depolarization of the post-synaptic calyx, and a resistive coupling for fast transmission between the Type I hair cell and post-synaptic calyx (reviewed in Contini et al., 2022 and modeled in Govindaraju et al. 2023).

As the *Vglut3-ko* model is missing the glutamatergic component of this complex input, our results raise the intriguing possibility that non-quantal input (the most likely alternate input) provided by the calyx environment could be driving the normal development of vestibular neurons. Perhaps it is not surprising then that calretinin and KCNQ4 expression develop normally in the central epithelial zones where non-quantal transmission has been experimentally observed (Songer and Eatock, 2013). If a large component of signaling between central zone hair cells and neurons is preserved, this may also explain why the knockout mice are able to traverse a balance beam without significant difficulty since those tests are effective at detecting central zone deficits (Ono et al., 2020). Whether there is significant non-quantal transmission in the peripheral/extrastriolar zones of vestibular epithelia remains to be determined. If all Type I hair cells are capable of non-quantal transmission, and since pure bouton-bearing vestibular neurons receiving input only from Type II hair cells are a very small minority of the total vestibular afferent population (Goldberg, 2000), then it is possible that most vestibular afferents receive a mixture of both quantal and non-quantal inputs. It would be useful if future work identified if or how the balance between the two modes of transmission varies by sensory context and epithelial zone.

VGN cell size is known to be correlated to zone of innervation; with the large calretinin positive soma belonging to the irregularly-firing calyx-only fibers innervating central epithelial zones (Kevetter & Leonard, 2002). In rat, large-diameter VGN have predominantly transient-firing patterns, consistent with the idea that these soma belong to the irregularly-firing group innervating the central zones (Kalluri et al., 2010; Kalluri, 2021; Bronson and Kalluri, 2023). Also in line with our previous findings in rat, here we found in both wildtype and Vglut3 KO mice that the distribution of VGN somatic size is bimodal with the same approximate range. Thus, the distribution of cell sizes is not impacted by the reduced input in the Vglut3 KO mice and cell size can serve as a rough marker for zone of innervation in both mouse models. Our finding that the large ganglion cells in the KO mice (as compared to in the wildtype mice) were more excitable with smaller net resting conductance indicates that these large VGN maybe compensating for a weaker sensory input by maintaining ion channel properties that make them more excitable. So, although the main thrust of this study suggests that the vestibular epithelium develops and functions normally in the Vglut3 KO mice, there are subtle changes in the cellular physiology consistent with a compensatory increase in excitability. However, this conclusion must be tempered given the sparsity of data in the large diameter VGN, especially at the oldest ages.

Finally, we acknowledge that the vestibular system works by integrating multiple sensory systems. This redundance means that even striking deficits in peripheral sensory signaling can produce minimal apparent balance disfunctions. Finding a functional test that accurately interrogates a specific deficit is a particular challenge for vestibular studies. In our case, the balance beam test is limited in scope and these animals may yet have vestibular deficits that went undetected.

Overall, the absence of glutamatergic input is a dramatic reduction of signaling from the inner hair cell to spiral ganglion neurons with a resulting significant impact on the diversification of those neurons. By contrast, the input signals driving vestibular neurons maybe more complex with both quantal glutamatergic and other (non-quantal or still other) mechanisms contributing to the net input. Definitive tests of the central hypothesis (that hair cell inputs drive neuronal diversity in the vestibular system) requires a more complete removal of hair cell input. This would entail blocking quantal and non-quantal transmission, as well as any tonic post-synaptic depolarization that may be caused by potassium accumulation in the calyx cleft. Alternatively, neuronal diversity in the vestibular epithelium maybe hardwired and not dependent on glutamatergic input.

### Funding

This work was supported by National Institutes of Health/National Institute on Deafness and Other Communication Disorders Grants R01 DC015512 to R.K., T32 DC009975 to K.R., and F32 DC020385 to D.B., as well as by the USC Provost’s Undergraduate Research Fellowship to R.C..

## Acknowledgments

We thank Dr. John Oghalai for providing the *Vglut3-ko* mice. We also thank Dr. Ksenia Gnedeva and Dr. Larry Hoffman for their comments on this work. Finally, we thank Dr. Seth Ruffins at the USC Optical Imaging Facility for his help in obtaining confocal images.

